# Probing Molecular Diversity and Ultrastructure of Brain Cells with Fluorescent Aptamers

**DOI:** 10.1101/2023.09.18.558240

**Authors:** Xiaotang Lu, Yuelong Wu, Peter H. Li, Tao Fang, Richard L. Schalek, Yaxin Su, Jeffery D. Carter, Shashi Gupta, Viren Jain, Nebojsa Janjic, Jeff W. Lichtman

## Abstract

Detergent-free immunolabeling has been proven feasible for correlated light and electron microscopy, but its application is restricted by the availability of suitable affinity reagents. Here we introduce CAptVE, a method using slow off-rate modified aptamers for cell fluorescence labeling on ultrastructurally reconstructable electron micrographs. CAptVE provides labeling for a wide range of biomarkers, offering a pathway to integrate molecular analysis into recent approaches to delineate neural circuits via connectomics.

## Main text

The brain is composed of a network of massively interconnected neurons, which collectively support a wide range of brain functions. In connectomics, automated serial section electron microscopy (ssEM) is used to reconstruct neural circuits at the synaptic level, in order to analyze the architectural organization and connectivity of the brain. However, the extraordinary diversity of neurons presents significant challenges for connectomic analysis – inferring the specific molecular types of neurons from EM is problematic. Without such knowledge, it would be difficult to determine the role of individual neurons in a circuit, and even more difficult to interpret the working principles of EM-reconstructed neural networks.

Correlated light and electron microscopy (CLEM) that superimposes immunofluorescence onto EM images^1^ can be used to directly identify cell types in EM image stacks^2^ or train deep learning algorithms to predict cell/synapse types^3^. Typically, immunohistochemistry requires cell permeabilization, a process using detergent to dissolve membranes to facilitate antibodies entering cells and accessing intracellular epitopes. Unfortunately, this process disrupts tissue ultrastructure, undermining neural circuit tracing. Prior research revealed that small immunolabels, like nanobodies^2^ and single-chain variable fragment antibodies^4^, can permeate cell membranes without the need for detergents. When extracellular space (ECS) is preserved, full-size antibodies can also be used for detergent-free immunolabeling^5^, although with reduced diffusion and restricted fine structure labeling^6^. As an increasing number of novel biomarkers are being discovered by single-cell transcriptomics^7,8^, the challenge now lies in expanding the array of labeling probes for permeabilization-free CLEM to improve the connection between ultrastructural analysis and the rapidly burgeoning knowledge of molecular cell diversity.

To meet this challenge, we introduce CAptVE (<u>C</u>orrelated <u>Apt</u>amer-assisted <u>V</u>olumetric <u>E</u>M), a method using slow off-rate modified aptamers (i.e., SOMAmer reagents) to probe the molecular signatures of individual brain cells within their ultrastructurally reconstructable connectome. We selected SOMAmer reagents for two main reasons: 1) Their specificities and affinity, thanks to the modification of the deoxyuridine base with hydrophobic residues^9^, are comparable to immunoglobulins for immunolabeling^10,11^; 2) The 50-nucleotide aptamers have molar mass (∼16 kDa) close to nanobodies (∼15 kDa), suggesting they might be applicable for detergent-free staining.

To examine the feasibility of using aptamers for detergent-free cell labeling, we applied the Cy5-conjugated anti-GFP SOMAmer reagents to paraformaldehyde-fixed mouse brains that express EGFP in a subset of GABAergic cortical interneurons (i.e., GIN mice). The tissues were prepared with ECS preservation but without detergent treatment. Since aptamers, particularly SOMAmer reagents, are selected for native unfixed protein targets, typical fixation conditions used for immunostaining might potentially decrease labeling efficiency by cross-linking proteins. Therefore, we began with mild fixation conditions^11^, using 2% paraformaldehyde at 4°C for 3 hours. Despite the colocalization of fluorescence, this mild fixation failed to preserve tissue integrity through the 2-day staining and imaging period, as demonstrated by the vacuoles on the EM image (Figure 1a-b). Consequently, we increased the paraformaldehyde concentration to 4%, maintaining the fixation conditions. GFP-SOMAmers bound correctly to EGFP, and the ultrastructure was well preserved (Figure 1c-d). Nevertheless, prolonged fixation, (e.g., 2% paraformaldehyde overnight, Figure S1), reduced the staining, probably owing to greater protein conformational changes and cross-linking.

**Figure 1.**
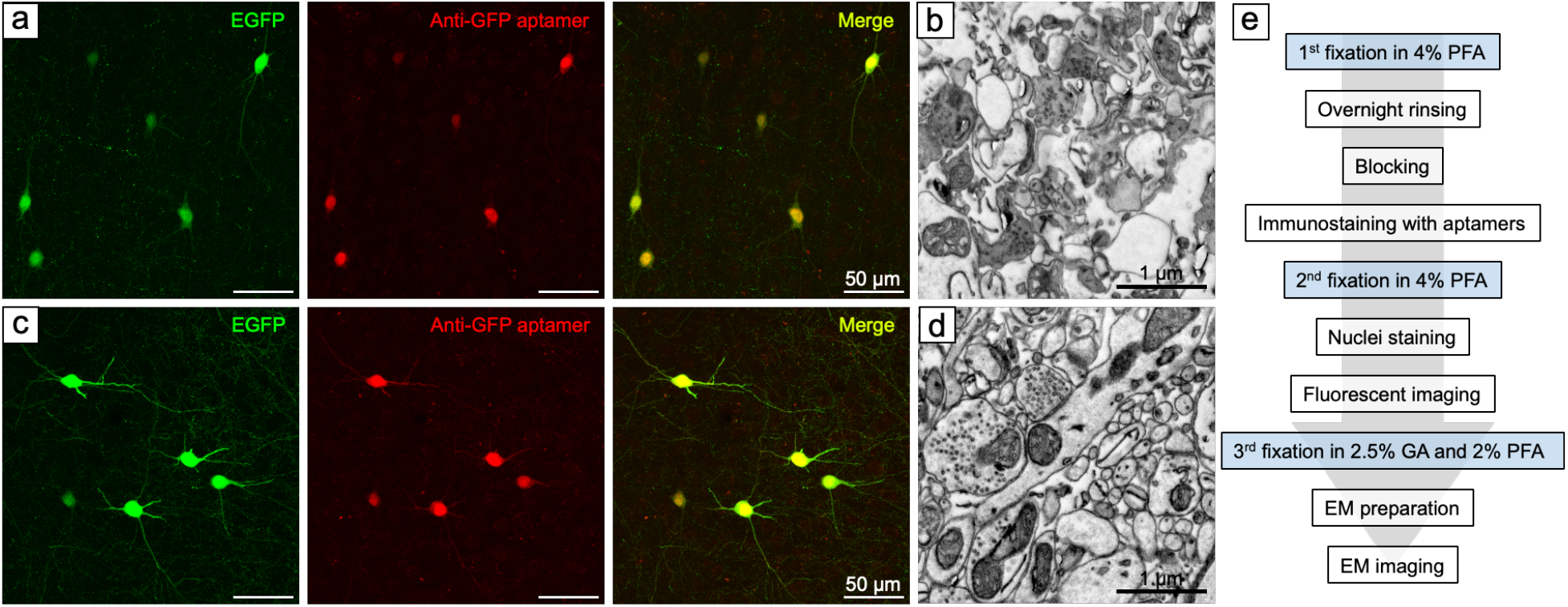
Impact of fixation on immunostaining and tissue ultrastructure. (a) GIN mouse cortex, initially fixed with 2% paraformaldehyde for 3 hours and stained with Cy5-conjugated GFP-SOMAmers, showed specific labeling. (b) This fixation was insufficient to preserve tissue ultrastructure. (c) Tissue fixed with 4% paraformaldehyde for 3 hours also showed specific labeling, and (d) the tissue ultrastructure was well preserved. (e) Overview of CAptVE workflow, highlighted three fixations.

We next investigated the potential of aptamers for volumetric CLEM using anti-calretinin (CR) and anti-neuropeptide Y (NPY) SOMAmer reagents. These biomarkers were chosen because they express in subtypes of cortical somatostatin (SST) interneurons^12^, whose ultrastructural differences are unknown. We first validated the anti-CR and anti-NPY SOMAmer reagents by comparing their fluorescence patterns with corresponding commercial antibodies (Figure 2a-b). Both reagents demonstrated fluorescence in the same cell populations as their corresponding antibodies. We then applied Cy5-conjugated anti-CR and Cy3-conjugated anti-NPY SOMAmer reagents in transgenic mice where GFP tagged the nuclear membranes of SST-positive interneurons (i.e., SST-cre;INTACT mice). The fluorescence-identified cells represented all seven possible labeling combinations of the three aptamers (in order of abundance: SST+/CR-/NPY-, SST-/CR+/NPY-, SST-/CR-/NPY+, SST+/CR-/NPY+, SST+/CR+/NPY-, SST+/CR+/NPY+ and SST-/CR+/NPY+). An analysis of fluorescent image stacks from the mouse somatosensory cortex (n=4, Figure S2-S5) revealed layer-specific abundance of these subtypes. In L1, SST-/CR-/NPY+ was the most abundant; SST-/CR+/NPY-predominated in L2/3 and L4, while SST+/CR-/NPY-was most abundant in L5 and L6 (Figure 2c).

**Figure 2.**
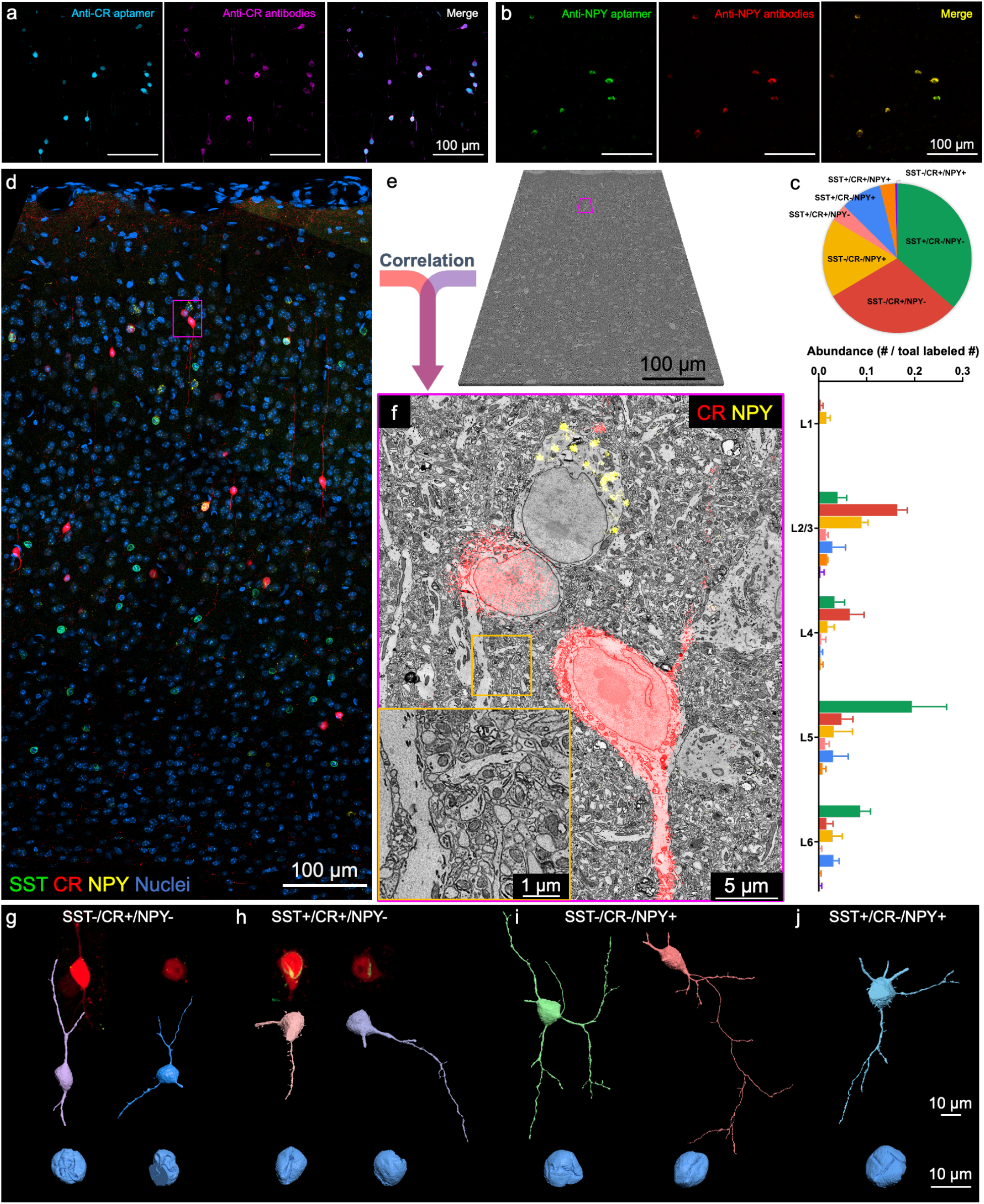
CAptVE demonstration. Comparison of (a) Calretinin (CR) and (b) Neuropeptide Y (NPY) SOMAmer reagents with corresponding commercial antibodies. (c) Abundance (top) and layer distribution (bottom) of the seven neuron subtypes expressing various combinations of SST, CR, and NPY. (d) Maximum intensity projection of the fluorescent LM stack and (e) the correlated EM stack. (f) Magnified view of the region outlined in magenta in (e and d), with superimposed fluorescent labels; the inset showed well-preserved and stained ultrastructure. (g-j) EM reconstructions of selected labeled cells and their nuclei. Additional reconstructions are available in Figure S6. Atop the SST-/CR+/NPY- and SST+/CR+/NPY-neuron reconstructions, fluorescent images displayed varying levels of CR expression.

In preparing the CLEM dataset, we stained and processed the tissue for EM imaging after confocal imaging, then co-registered the two image stacks manually (Figure 2d-f). The EM reconstructions allowed the assessment of morphological and ultrastructural features of the labeled cells. The six most common cell expression patterns were found in the EM volume, but the rare SST-/CR+/NPY+ type (i.e., 2 out of 451 labeled cells) was not present. The CR-only (SST-/CR+/NPY-) neurons exhibited more nuclear infolding with an average surface-area-to-volume ratio of 1.57 (n=3) compared to the other cell types whose ratios ranged from 0.97 to 1.26 (n=9). These CR-only neurons however were further divisible into two morphological types: bipolar and multipolar. Moreover, the bipolar CR-only neurons exhibited stronger red fluorescence than their multipolar counterparts, further arguing that these are two distinct classes (Figure 2g). Calretinin fluorescence in SST+/CR+/NPY-neurons also corresponded to cells with distinct morphologies: those with stronger CR fluorescence were spiny, while those with lower expression had smoother dendrites (Figure 2h). We also observed diverse morphologies within other cell types. For example, NPY-only (SST-/CR-/NPY+) neurons exhibited two different somatic morphologies — spindle-shaped and polygonal (Figure 2i and Figure S6). Even the cells labeled for the three biomarkers (i.e., SST+/CR+/NPY+), were variable in shape (Figure S6). This analysis underscores the challenges in defining a cell type by sparse molecular categories and varying protein expression levels. Future efforts will need to increase the multiplexity of labeling in order to refine cell type analysis.

Lastly, we demonstrated that aptamers could reveal both previously identified and newly discovered biomarkers in brain cells. We leveraged the SOMAmer library, which contains over 7000 validated reagents for proteomic assays. However, considering those reagents were originally selected for native unfixed human proteins, we began by examining their ability to bind epitopes in fixed mouse brain samples. We performed full-panel SOMAscan assays, which measured the concentrations of the proteins bound by SOMAmer reagents, on both frozen and paraformaldehyde-fixed mouse brain tissues. We found that 1132 SOMAmer reagents specifically bound to their protein targets in frozen mouse brains (i.e., we used a binding threshold of 3× background -see Figure S2). Post paraformaldehyde treatment, the binding affinity decreased for 95.2% of the reagents, confirming the negative impact of fixation on immunolabeling. Still, 481 reagents maintained sufficient binding in fixed mouse brain tissues.

Drawing on several resources, ^13–17^ we identified a set of biomarkers from the 481 SOMAmer reagents that were likely expressed only in subsets of brain cells. Our results confirmed that SOMAmer reagents effectively bind to a range of target proteins by demonstrating that individual probes label distinct cell types in various brain regions (Figure 3). For example, we found that peroxiredoxin 6 (PRDX6) reagents specifically labeled the somata and endfeet of astrocytes that ensheath mouse cerebral cortex vasculature, in contrast to another astrocyte marker, glial fibrillary acidic protein (GFAP), which primarily localizes to astrocytic processes. Quinoid dihydropteridine reductase protein (QDPR) reagents marked a subset of oligodendrocytes, while myelin associated glycoprotein (MAG) was limited to the myelin sheath. In terms of neurons, in addition to canonical interneuron markers such as CR and NPY (see above), we labeled cells expressing proenkephalin (PENK) which is a newly discovered interneuron marker, and cells expressing neurogranin (NEUG), which is a marker for pyramidal neurons. We also found protein kinase C gamma (PRKCG) labeled Purkinje cells in the cerebellum as well as a subset of L2/3 and L5 neurons in the somatosensory cortex. In the hippocampus, ELAV like RNA binding protein 2 (ELAVL2) labeled hilar interneurons in the dentate gyrus and CA3 pyramidal neurons. This protein was also highly expressed in L5 neurons of the somatosensory cortex. Neuronal pentraxin receptor (NPTXR) reagents labeled subsets of neurons in the granule layer of the dentate gyrus and their processes in the hilus. This aptamer also labeled subgroups of cortical neurons and their processes.

**Figure 3.**
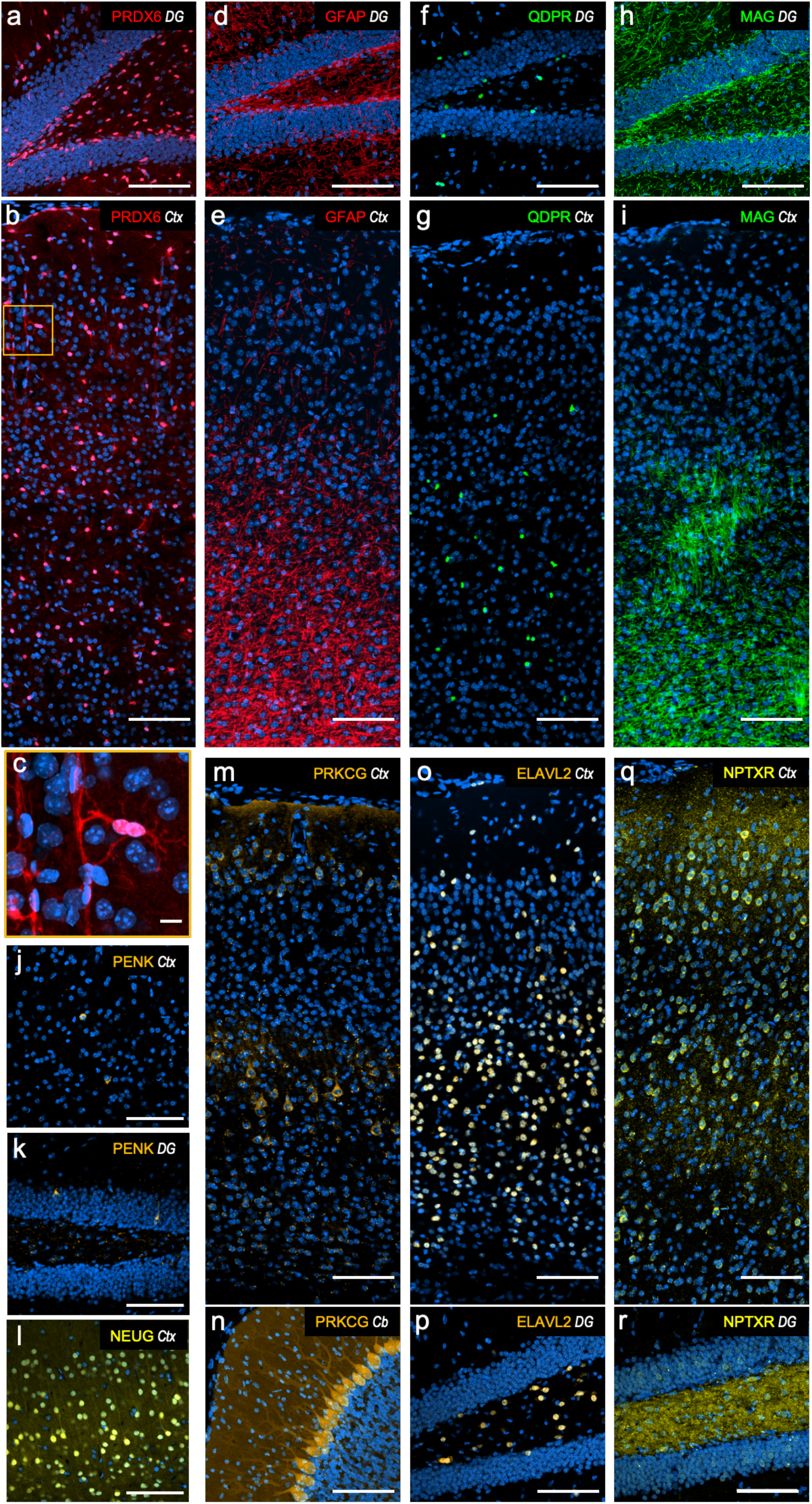
Detergent-free immunostaining with selected SOMAmer reagents in various brain regions (Ctx: Cortex; DG: Dentate Gyrus; Cb: Cerebellum). The scale bar in (c) is 10 μm, and in all other panels are 100 μm.

In summary, we have demonstrated the use of fluorescently tagged SOMAmer reagents for high-quality volumetric CLEM. By superimposing multiple fluorescent labels on an EM image stack, we were able to distinguish various cell types and then seek the ultrastructural properties of the same cells. We also showed that the novel biomarkers identified by single-cell transcriptomics can be easily adapted by SOMAmer reagents for proteomic cell labeling. We now plan to expand the aptamer library to include biomarkers verified by recent transcriptomic studies. Considering the ease of generating enormously large libraries of randomized DNA sequences for efficient in vitro selection, we believe that CAptVE will allow us to connect the collective knowledge of brain cell types with connectomic analysis, in order to routinely integrate ultrastructure and connectivity analysis into a comprehensive understanding of brain cell types and their role in neuronal circuitry.

## Online Methods

### Animals

Adult GIN mice (FVB-Tg(GadGFP)45704Swn/J, JAX strain #:003718) were used to develop the fixation and staining protocol with anti-GFP SOMAmer reagents. Adult C57BL/6 mice (JAX strain #: 000664) were used to test other SOMAmer reagents. Adult SST-cre;INTACT mice, offsprings of SST-IRES-cre (JAX strain #: 013044) and INTACT mice (JAX strain #: 021039), were used to acquire the CLEM datasets. All animal procedures were performed according to US National Institutes of Health guidelines and approved by the Committee on Animal Care at Harvard University.

### Materials

NaCl (S3014), NaHCO3 (S5761), NaH2PO4·H2O (567549), KCl (P9541), glucose (G7021), PBS powder (P3813), mannitol (M4125), 1 M CaCl2 solution (21115), 1 M MgCl2 (M1028), 1 M MgSO4 solution (M3409), dextran sulfate sodium salts (m.w. 6500-10000, D4911), potassium ferrocyanide (60279) were purchased from Sigma-Aldrich.

Glycine (J16407), sheared salmon sperm DNA (10 mg/mL, AM9680), bovine serum albumin (50 mg/mL, AM2616), 1 M HEPES solution (15630080), 0.5 M EDTA solution (15575020), Hoechst 33342 Solution (62249), normal goat serum (PCN5000), 120 μm deep Secure-Seal spacer (S24737), T-PER tissue protein extraction reagent (78510), Halt protease inhibitor cocktail (87785), Micro BCA protein assay kit (23235) were purchased from ThermoFisher.

Sodium cacodylate trihydrate (12310), 32% paraformaldehyde solution (15714), 25% glutaraldehyde solution (16220), 4% OsO4 solution (19190), TCH (21900), and acetonitrile (10020) were purchased from Electron Microscopy Sciences.

LX112 (21310), NSA (21355), NMA (21350), BDMA (21365) were purchased from Ladd. Anti-Calretinin antibodies (rabbit, C7479) were purchased from Sigma-Aldrich. Anti-NPY antibodies (rabbit, 11976) were purchased from Cell Signaling Technology. Alexa594 Fab2 fragment antibodies (donkey-anti-rabbit, 711-586-152) were purchased from Jackson ImmunoResearch.

Polyanionic competitor (“Z-block”, 30-mer DNA that contains modified nucleotides Bn-dU^10^ with the following sequences: 5’-[AC[Bn-dU]2]7AC-3’)^11^ was supplied by SomaLogic.

### Solutions

The recipes for the solutions used in sample preparation are as follows.

- aCSF: 125 mM NaCl, 26 mM NaHCO3, 1.25 mM NaH2PO4, 2.5 mM KCl, 20 mM glucose, 1 mM MgCl2, and 2 mM CaCl2. MgCl2 and CaCl2 were added after the solution was bubbled with carbogen gas for 20 min.
- LM fixative: 1× PBS, 4% w/v mannitol, 4% w/v PFA.
- Aptamer dilution buffer: 5 mM HEPES, 1 mM EDTA (pH 7.5).
- Aptamer blocking buffer: 1× PBS, 5 mM MgCl2, 100 mM glycine, 5 mg/mL BSA, 0.5 mg/mL salmon sperm DNA.
- Aptamer immunostaining buffer: 1× PBS, 5 mM MgCl2, 1 mM dextran sulfate (assuming average m.w. 8000), 10 mg/mL BSA, 0.5 mg/mL salmon sperm DNA, 25 μM polyanionic competitor (“Z-block”).
- Antibody blocking buffer: 1× PBS, 10% normal goat serum, 100 mM glycine, 0.05% NaN3.
- Antibody immunostaining buffer: 1× PBS, 3% normal goat serum, 100 mM glycine, 0.05% NaN3.
- EM fixative: 150 mM sodium cacodylate solution (pH 7.4), 4 mM MgSO4, 2 mM CaCl2, 4 w/v% mannitol, 2 w/v% PFA, 2.5 w/v% glutaraldehyde.
- LX112 resin: 15.6 g LX112, 4.7 g NSA, 9.7 g NMA, 0.6 g BDMA. The resin should be well mixed and degassed before use.

### Perfusion and tissue fixation

Mice were perfused and fixed using the ECS-preserving protocol (CITE). Briefly, deeply anesthetized mice were transcardially perfused at a flow rate of 10 mL/min with the following solutions: 1) aCSF for 2-3 minutes, 2) 15 w/v% mannitol dissolved in aCSF for 1 minute, 3) 4.5 w/v% mannitol dissolved in aCSF for 5 minutes, 4) ice-cold LM fixative for 5 minutes. After perfusion, the brains were immersed in LM fixative for 2 hours with gentle agitation at 4°C. The brains were sectioned into 120 μm slices using a Leica VT1000 S vibrating blade microtome in a cold solution containing 4% w/v mannitol and 1x PBS. Slices were post-fixed in cold LM fixative for 1 hour (total fixation time 3 hours). The fixed slices were then rinsed with the cold solution containing 4% w/v mannitol and 1x PBS for 3 x 5 min and stored in the same solution at 4°C for up to one month.

### Sample preparation for SOMAscan assays

Native mouse brain tissue was acquired by blood removal with aCSF, cutting 200 μm sagittal sections near the midline in cold aCSF, and snap freezing on dry ice before storage at -80°C. Fixed mouse brain tissue was fixed as described above and then snap frozen. For lysis, 1 μL Halt protease inhibitor cocktail was added to every 100 μL of T-PER tissue protein extraction reagent and 200 μL of this extraction buffer per 10 mg of tissue was used for lysis using a Qiagen TissueLyser II. The lysate was centrifuged at 14,000 x g for 10 min at 4°C, the supernatant was collected, and protein concentration was determined using a microBCA protein assay kit. Samples were then normalized to 75 μL at 200 μg/mL total protein concentration using PBS before the assay.

### Reagent storage

SOMAmer reagents were diluted in the reagent dilution buffer to create 10 μM aliquots, which were stored at –20°C. Working aliquots were moved to a 4°C refrigerator. Before use, the reagents were subjected to a heat-cooling process (90°C for 5 min, then cooled down on ice) and mixed with the immunostaining buffer to achieve a final concentration of 100 nM.

### Aptamer-based immunostaining and fluorescent imaging

Except for fluorescent imaging, all procedures were conducted at 4°C to maintain membrane integrity. The fixed 120 μm brain slices stored in the mannitol-added PBS solution were transferred to a petri dish. The region of interest was dissected from the brain slices and transferred to a 48-well plate. After 3 × 5 min rinse in a solution containing 4% w/v mannitol and 1× PBS, 250 μL of aptamer blocking buffer was added to each well. The plate was then placed on a nutating mixer for gentle agitation. The sections were blocked for 5 hours to overnight. Before the next step, 2.5 μL of the stock SOMAmer reagent (10 μM) was diluted in 250 μL of aptamer immunostaining buffer to make a final concentration of 100 nM. After blocking, the immunostaining solution was added to the well. After 2 days of staining, the samples were rinsed with PBS for 3 × 10 min. For CLEM, the sample needs to be post-fixed in LM fixative for 12-24 hours. After post-fixation, the samples were rinsed with PBS for 3 × 10 min and stained overnight in a diluted Hoechst nuclei staining solution (1:5000 in PBS) overnight. For an LM-only experiment, post-fixation is not needed. The samples were stained with diluted Hoechst solution directly after rinsing. The next day, the samples were rinsed with PBS for 3 × 10 min, then mounted onto glass slides with a 120 μm Secure-Seal spacer for fluorescent imaging. A Zeiss LSM 900 confocal laser scanning microscope equipped with a 20x/0.8 NA air-objective was used for fluorescent imaging.

### Co-immunostaining with full-size antibodies for aptamer validation

All procedures were conducted at 4°C unless otherwise specified. After aptamer staining, the samples were post-fixed in LM fixative for 12-24 hours to fortify them for subsequent staining. The sample were then rinsed in PBS for 3 × 10 min and blocked in antibody blocking buffer for 5 hours. Meanwhile, the primary antibody immunostaining solution was prepared by diluting the antibody stock with the antibody immunostaining buffer at a ratio of 1:100. After blocking, the primary antibody immunostaining solution was added to the well-plate and incubated for 3-5 days. The samples were then rinsed with PBS for 3 × 10 min and left in PBS overnight. The next day, the fluorescent secondary antibody immunostaining solution (prepared in the same manner as the primary antibody) was added to the samples and incubated for 3 days. After staining, the samples were rinsed with PBS for 3 × 10 min and stained overnight in diluted Hoechst nuclei staining solution (1:5000 in PBS) overnight. The next day, the samples were rinsed with PBS for 3 × 10 min, then mounted onto glass slides with a 120 μm Secure-Seal spacer for fluorescent imaging.

### EM sample preparation

After fluorescent imaging, tissues were post-fixed in EM fixative at 4°C for a minimum of one day before EM staining. A modified ROTO protocol was used to stain tissues for EM imaging. All procedures were conducted at room temperature unless specified otherwise. Samples were rinsed 3 × 10 min in a 150 mM sodium cacodylate buffer (pH 7.4) and stained on a rotator for 1 hour with reduced-osmium solution containing 1 w/v% OsO4, 1.5 w/v% potassium ferrocyanide, and 150 mM cacodylate buffer (pH 7.4). After the first osmication, samples were rinsed 3 × 10 min in ddH2O and stained in filtered 0.5 w/v% thiocarbohydrazide aqueous solution for 30 min. Then samples were rinsed 3 × 10 min in ddH2O and stained with 2 w/v% OsO4 aqueous solution for 1 hour. After 3 × 10 min rinses, samples were transferred to 1 w/v% uranyl acetate aqueous solution, wrapped in foil to protect from light, for overnight staining (∼12 hours). The next day, samples were washed 3 × 10 min in ddH2O. Stained samples were dehydrated by passing through a graded series of acetonitrile (25, 50, 75, 100, 100%, 10 min each). After dehydration, samples were infiltrated at room temperature with 25% LX112 resin:acetonitrile for 1 hour, 50% resin:acetonitrile for 2 hours, 75% resin:acetonitrile for 3 hours, 100% fresh resin overnight, 100% fresh resin for 8-12 hours on a rotator. Fully infiltrated samples were placed in a flat embedding mold and cured in a 60°C oven for two days.

### EM imaging and image processing

The resin-embedded sections were cut into serial 30 nm ultrathin sections and collected on carbon-coated Kapton tape using an automated tape-collecting ultramicrotome (ATUM) ^18^. The tape was cut into strips and mounted onto silicon wafers. Those sections were then post-stained with filtered 4 w/v% uranyl acetate aqueous solution for 4 minutes, followed by Leica Ultrostain II lead citrate solution for 4 minutes. We used Zeiss MultiSEM 505 scanning electron microscope with 61 electron beams to acquire 4 nm × 4 nm resolution EM images from the defined ROI. The raw EM images were stitched and aligned with a custom Python code, called FEABAS. Detailed instructions on installing and using FEABAS are available at https://github.com/YuelongWu/feabas.

### Automated segmentation and proofreading

The flood-filling networks model ^19^, previously trained on the human cortex H01 dataset ^20^ at 32 × 32 × 30 nm resolution, was employed to auto-segment our EM dataset (data is accessible via https://lichtman.rc.fas.harvard.edu/aptamer_clem/). Guided by the superimposed fluorescence image, we picked labeled neurons from the EM dataset. Base supervoxels from automatic segmentation were grouped into larger per-cell segments for these labeled neurons. We then manually proofread to refine the segmentation.

## Acknowledgments

We thank Prof. Larry Gold for his support of this collaborative project.

We thank the team at SomaLogic for synthesizing the SOMAmer reagents used in this project. SOMAmer reagent is a registered trademark of SomaLogic Operating Company, Inc.

We thank Dr. Simon T. Dillon and Prof. Towia Libermann for the help with SOMAscan assays. We thank Dr. Daniel Berger for his help with rendering 3D models.

Special thanks to Jingjing (Sherry) Wu, Shuhan Huang, and Prof. Gordon Fishell at Harvard Medical School for providing the transgenic SST-cre;INTACT mice and for their helpful discussions.

This research is supported by NIH BRAIN Initiative award K99MH128891 and NIH grant U19NS104653 and UG3MH123386.

## Author contributions

X.L. and T.F. conceived the idea. X.L. designed and implemented the experiments and analyzed the data with helpful inputs from all authors. J.D.C., S.G., and N.J. contributed to the generation of SOMAmer reagents and provided suggestions on their use. R.L.S. assisted in collecting serial ultrathin sections and EM imaging. Y.W. stitched and aligned EM images. P.H.L. conducted auto-segmentation. Y.S. proofread auto-segmentation and performed 3D rendering. X.L. and J.W.L. wrote the manuscript, and all authors reviewed and approved the manuscript.

## Supplementary information

**Figure S1.**
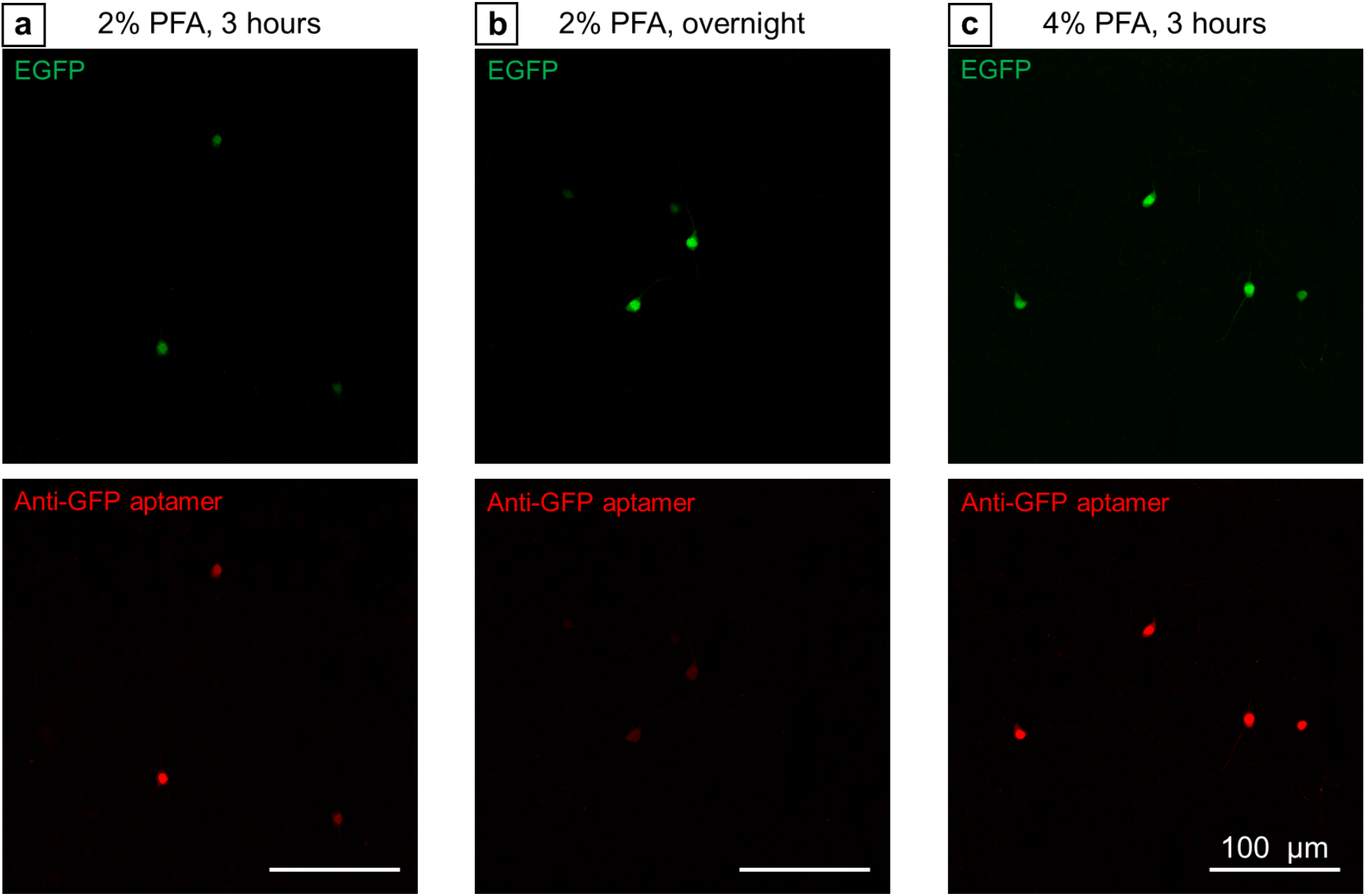
Impact of Fixation on EGFP Preservation and Immunostaining. (a) GIN mouse cortex fixed with 2% paraformaldehyde for 3 hours, stained with Cy5-conjugated GFP-SOMAmers. (b) Tissue fixed with 2% paraformaldehyde overnight, stained with SOMAmer reagents. (c) Tissue fixed with 4% paraformaldehyde for 3 hours, stained with SOMAmer reagents. The colocalization of red and green fluorescence in all cases signifies specific labeling. The decreased green fluorescence in (a) indicates poor EGFP preservation due to mild fixation. Overnight fixation in (b) results in well-preserved EGFP but decreased red fluorescence, suggesting possible conformational changes or excessive cross-linking affecting aptamer binding. Both strong EGFP preservation and aptamer labeling were achieved with 4% paraformaldehyde fixation for 3 hours in (c).

**Figure S2.**
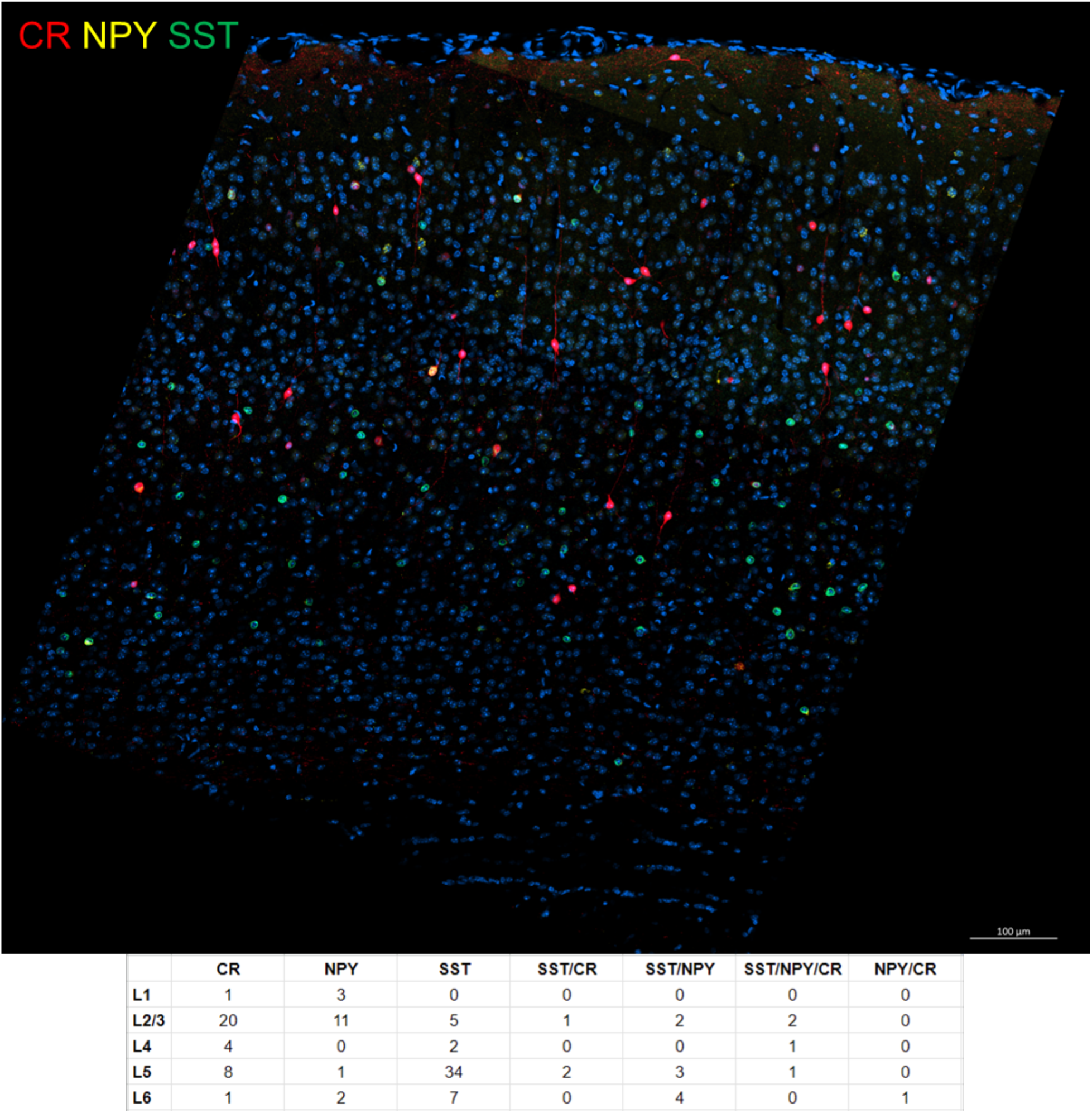
Fluorescent image stack #1 and cell type distribution. (Top) A max intensity projection of 10 μm optical slices from the image stack #1 of a transgenic SST-cre;INTACT mouse somatosensory cortex. Calretinin (red) and Neuropeptide Y (yellow) were labeled with SOMAmer reagents, while SST neuron nuclei were tagged with GFP (green). (Bottom) Distribution statistics of various labeled cell types across six cortical layers.

**Figure S3.**
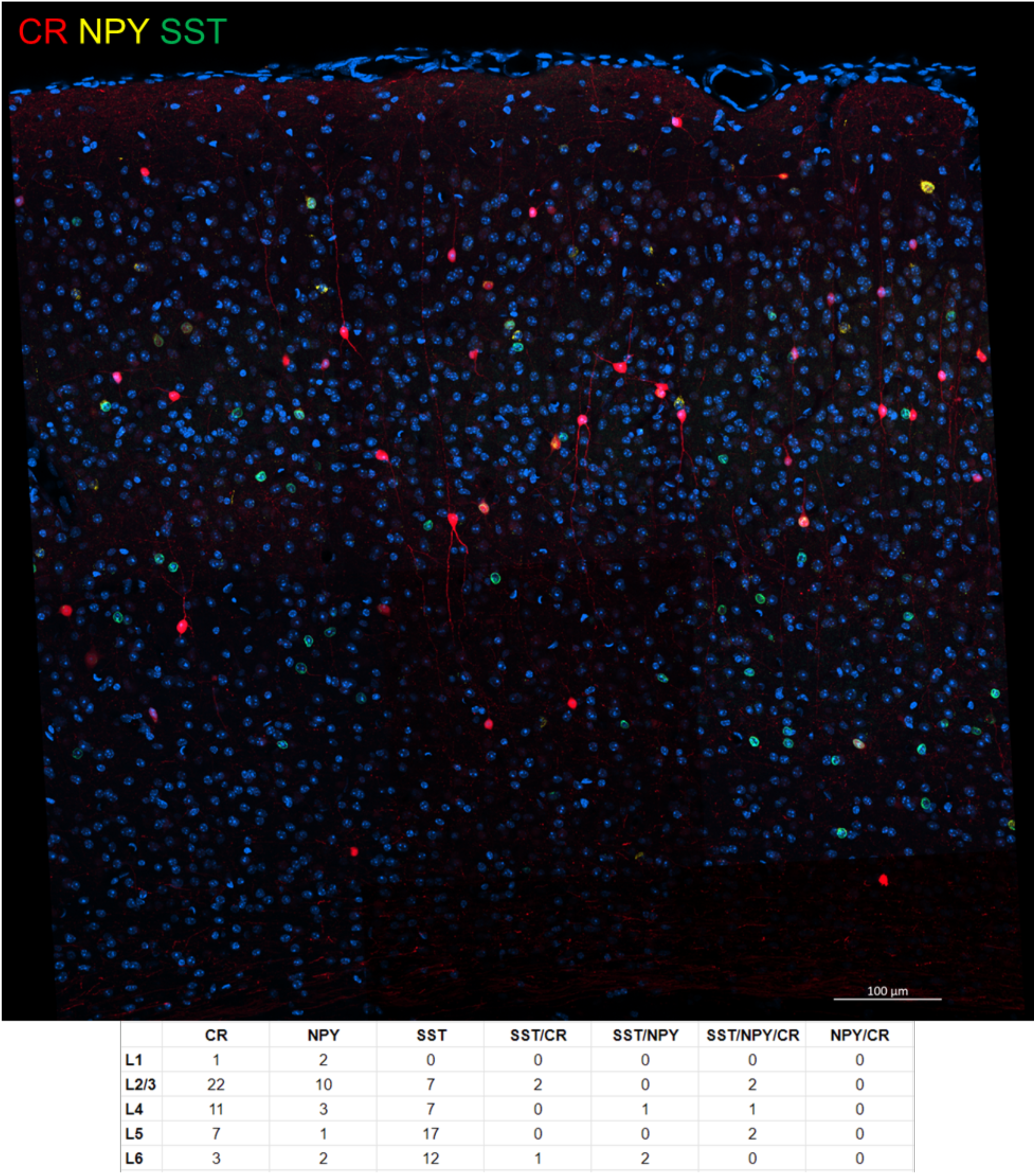
Fluorescent image stack #2 and cell type distribution. (Top) A max intensity projection of 10 μm optical slices from the image stack #2 of a transgenic SST-cre;INTACT mouse somatosensory cortex. Calretinin (red) and Neuropeptide Y (yellow) were labeled with SOMAmer reagents, while SST neuron nuclei were tagged with GFP (green). (Bottom) Distribution statistics of various labeled cell types across six cortical layers.

**Figure S4.**
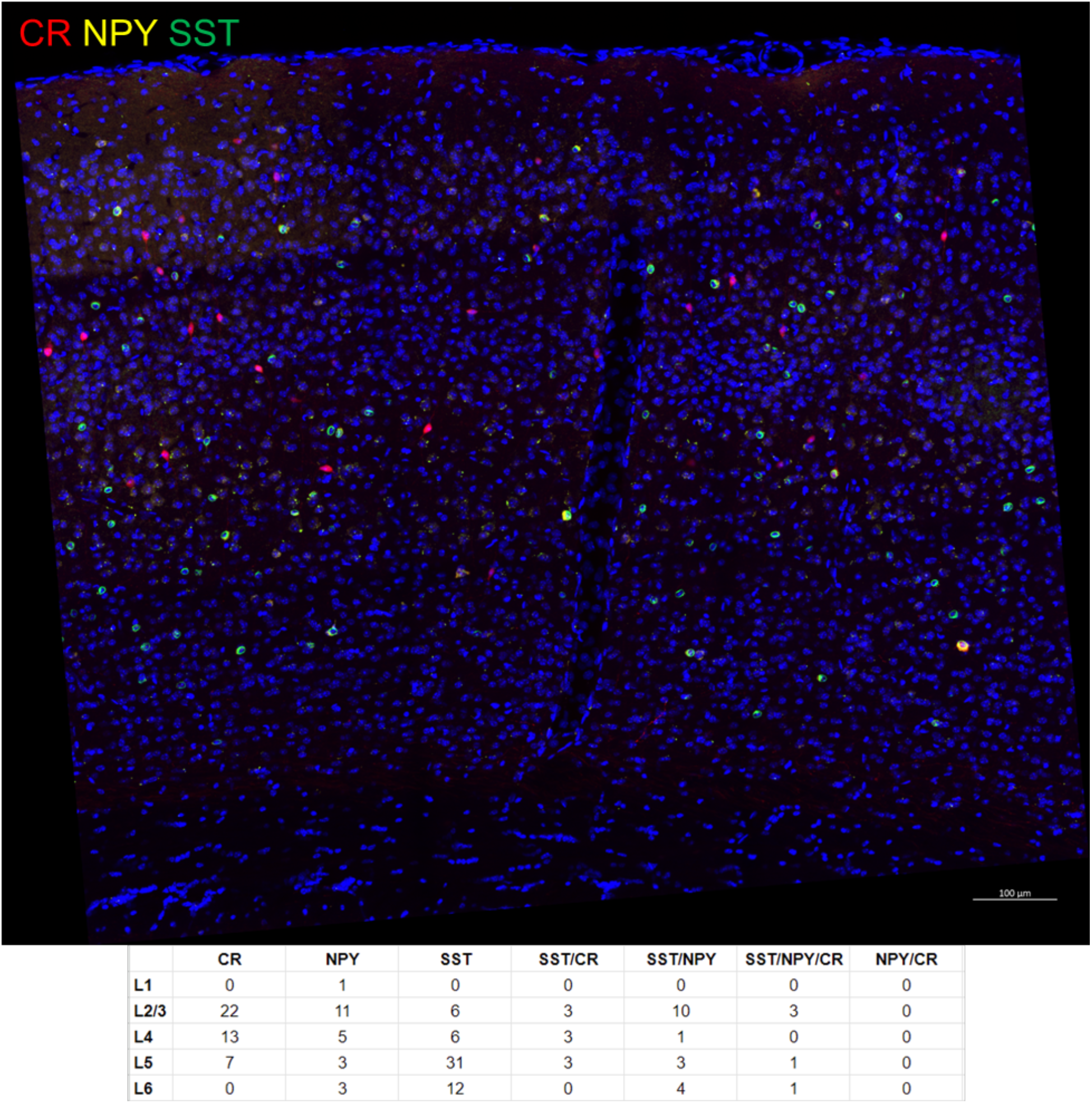
Fluorescent image stack #3 and cell type distribution. (Top) A max intensity projection of 10 μm optical slices from the image stack #3 of a transgenic SST-cre;INTACT a select few mouse somatosensory cortex. Calretinin (red) and Neuropeptide Y (yellow) were labeled with SOMAmer reagents, while SST neuron nuclei were tagged with GFP (green). (Bottom) Distribution statistics of various labeled cell types across six cortical layers.

**Figure S5.**
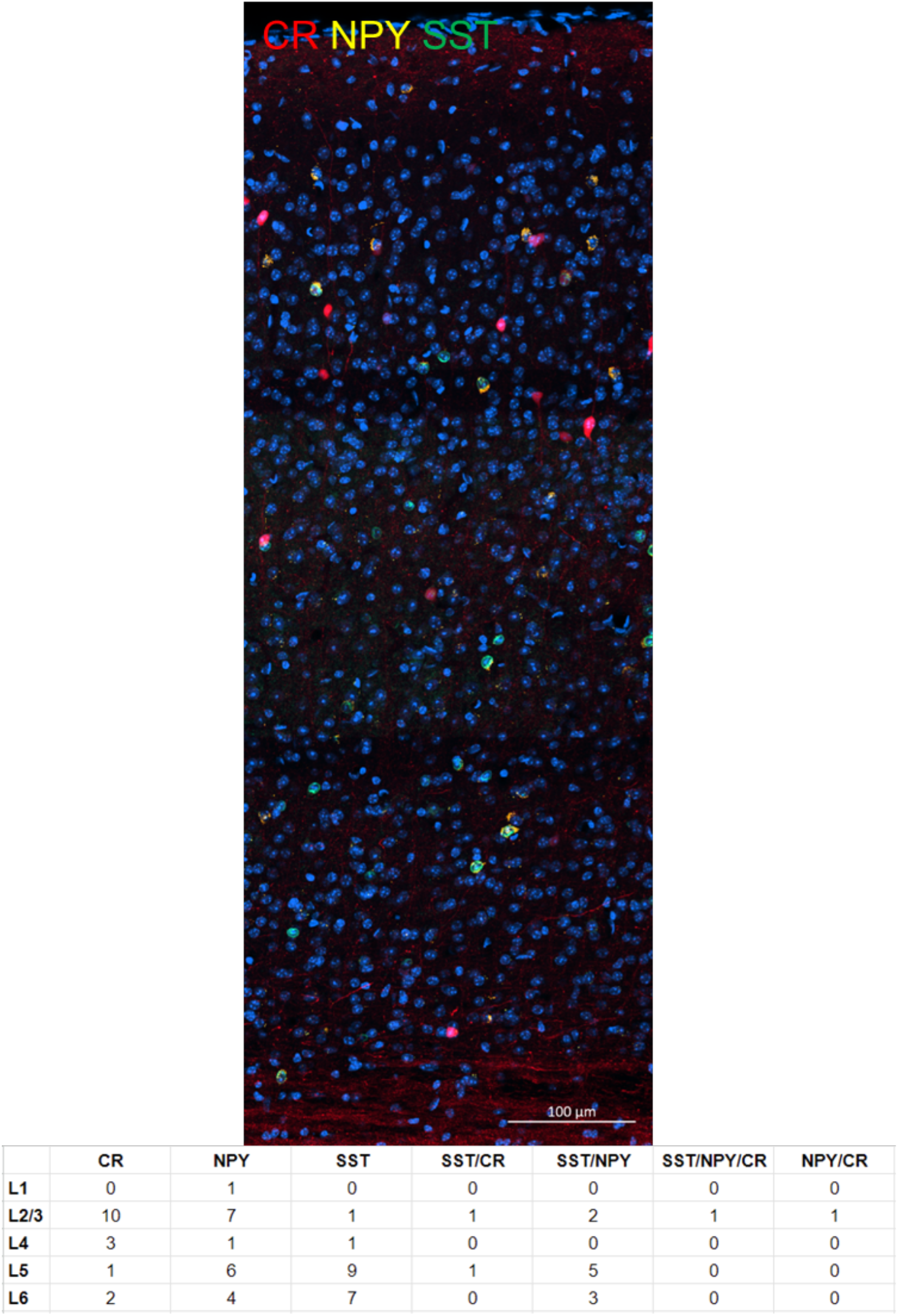
Fluorescent image stack #4 and cell type distribution. (Top) A max intensity projection of 10 μm optical slices from the image stack #4 of a transgenic SST-cre;INTACT mouse somatosensory cortex. Calretinin (red) and Neuropeptide Y (yellow) were labeled with SOMAmer reagents, while SST neuron nuclei were tagged with GFP (green). (Bottom) Distribution statistics of various labeled cell types across six cortical layers.

**Figure S6.**
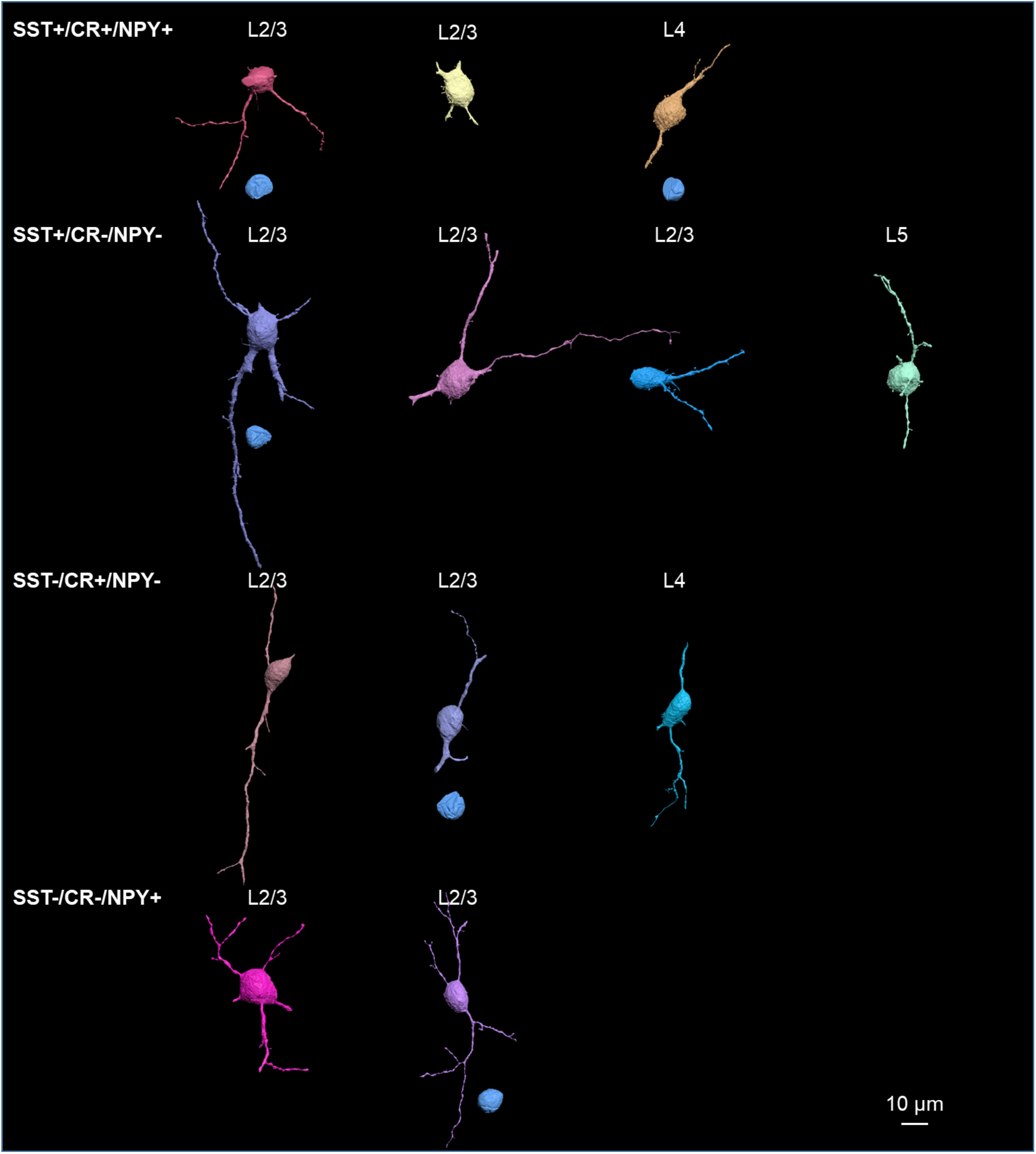
EM reconstructions of labeled cells and their nuclei that were not presented in Figure 2.

**Figure S7.**
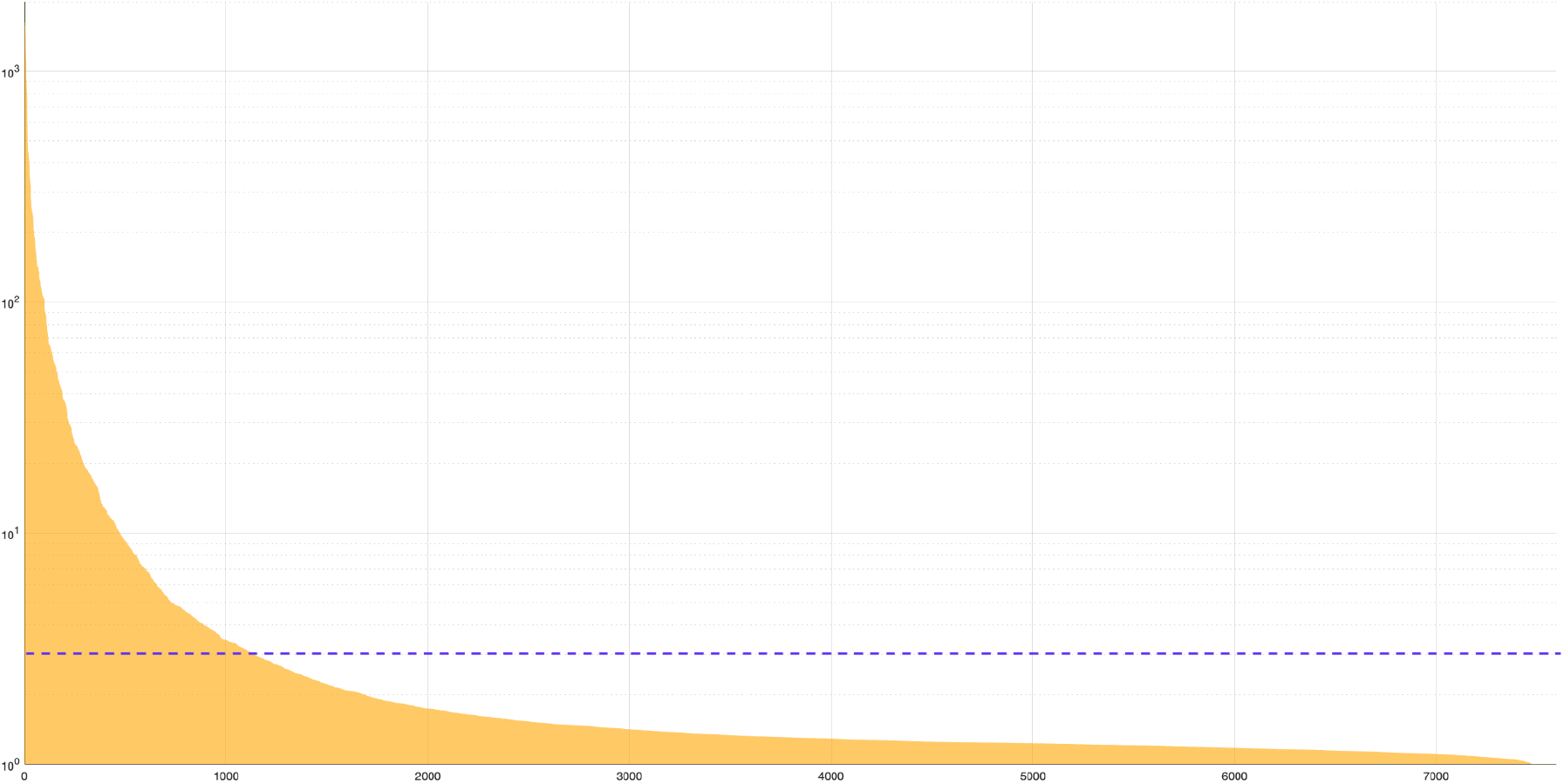
Analysis of SomaScan assay on a frozen mouse brain sample. The Relative Fluorescence Units (RFU) for each protein target were measured and normalized according to SomaLogic’s protocol. The normalized RFU for each protein target was then divided by the buffer RFU to derive a relative value. Protein targets were ranked based on these relative RFU values from high to low. The y-axis, plotted on a log scale for better visualization, represents these relative values. In this context, a value of 1 represents the background buffer binding. Each yellow line represents the relative binding of a protein target, while the purple dashed line marks the threshold of 3× buffer binding. In this analysis, we identified 1132 targets that exhibited binding above the purple dashed line, indicating significant binding by SOMAmer reagents. Due to space limitations on the x-axis, it was not possible to label all target names; instead, we only labeled the id numbers of the targets after sorting by RFU values.

**Figure S8.**
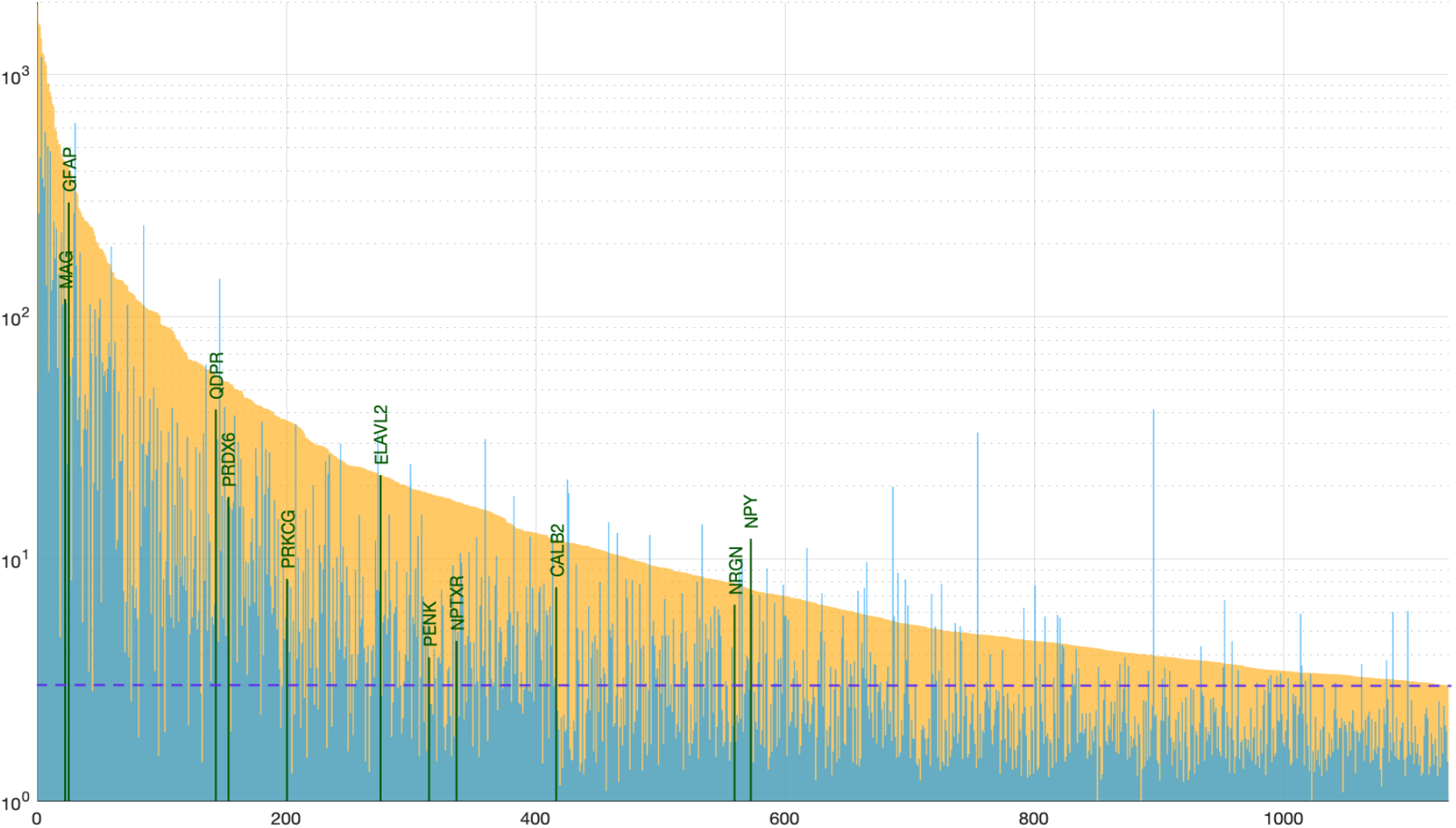
Comparative analysis of SomaScan assays on frozen and aldehyde-fixed mouse brains. From Figure S7, we selected 1132 protein targets that demonstrated 3× background buffer binding in the frozen mouse brain, represented by yellow lines. We then plotted the binding of these same targets in the aldehyde-fixed mouse brain, represented by blue lines, at the same position. This allowed direct visualization of changes in binding after fixation. The data revealed that the SOMAmer binding affinity decreased for 95.2% of the targets. Out of the 481 targets that maintained over 3× background binding after fixation (above the purple dash line), we highlighted the SOMAmer reagents selected for this study in dark green, with their corresponding gene names labeled on top.

